# High rates of spontaneous chromosomal duplications unravel dosage compensation by translational regulation

**DOI:** 10.1101/2022.02.03.478961

**Authors:** Marc Krasovec, Remy Merret, Frédéric Sanchez, Sophie Sanchez-Brosseau, Gwenaël Piganeau

## Abstract

While duplications have long been recognized as a fundamental process driving major evolutionary innovations, direct estimations of spontaneous chromosome duplication rates, leading to aneuploid karyotypes, are scarce. Here, we provide the first estimations of spontaneous chromosome duplication rates in six unicellular eukaryotic species from mutation accumulation (MA) experiments. The spontaneous chromosome duplication rates reach 1×10^−4^ to 1×10^−3^ per genome per generation, which is ~4 to ~50 times less frequent than spontaneous point mutations per genome, whereas chromosome duplication events can affect 1 to 7% of the total genome size. Comparative transcriptomics between MA lines with different chromosome duplications reveals a strong positive correlation between RNA expression rate and DNA copy number. However, comparative analyses of the translation rate of mRNAs estimated by polysome profiling unravel a chromosome specific dosage compensation mechanism. In particular, one chromosome with a gene average of 2.1 excess of mRNAs is compensated by an average of ~0.7 decrease in translation rates. Altogether, our results are consistent with previous observations of a chromosome dependent effect of dosage compensation and provide evidence that it may occur during translation. These results support the existence of a yet unknown post-transcriptional mechanism orchestrating the modification of translation of hundreds of transcripts from genes located on duplicated regions in eukaryotes.

## INTRODUCTION

Complete or partial chromosome duplications leading to aneuploidy karyotypes are known to be at the origin of genetic diseases, such as Trisomy 21 in humans, and is a near universal feature of tumor cells (Rajagopalan and Lengauer 2004). When a single chromosome is duplicated, an immediate issue arises: the imbalance of ploidy and gene dose within a same karyotype. This is expected to have deleterious effects because it creates an imbalance of the transcript and protein productions, which is costly and may disrupt the function of a pathway and protein interactions (Dephoure et al. 2014; Veitia and Potier 2015). The deleterious effects of aneuploidy have been documented in many different biological model systems such as yeast (Torres et al. 2007), mouse and human (Gearhart et al. 1987). In *Caenorhabditis elegans*, mutation accumulation experiments provided evidence of purifying selection against gene duplications causing excess of transcripts, as compared to gene duplications associated to invariant transcript levels (Konrad et al. 2018). Moreover, the consequences of aneuploidy on gene expression are complex and while gene expression may increase with chromosome copy number, gene expression changes may also spread outside the duplicated regions in *Arabidopsis* (Hou et al. 2018; Song et al. 2020), *Drosophila* (Devlin et al. 1988), and human cells (FitzPatrick et al. 2002). Contrasting with the before mentioned studies, aneuploidy may confer a selective advantage as a response to stress, such as resistance to drugs in *Candida albicans* (Selmecki et al. 2006) or *Saccharomyces cerevisiae* (Chen et al. 2012). Recent evidence about a lack of deleterious effect of aneuploidy is the high aneuploidy prevalence and tolerance across *Saccharomyces cerevisiae* lineages (Scopel et al. 2021), a fifth of the sequenced strains harboring atypical aneuploidy karyotypes (Peter et al. 2018). Indeed, different dosage compensation mechanisms have evolved and may restore the ancestral gene dose leading to aneuploidy tolerance. The three targets of dosage compensation mechanisms of duplicated genes are the modification of the transcription rate, the translation rate, or the protein degradation rate of duplicated genes. The most studied mechanisms are those involved in modifying the transcription rate during the evolution of heterogametic sex chromosomes, known both in plants (Muyle et al. 2012; Muyle et al. 2017; Charlesworth 2019) and animals (Disteche 2012; Graves 2016). Different mechanisms evolved either by simulating the ancestral ploidy by doubling the expression of the genes of the single copy male X (Baker et al. 1994); or by equalizing the ploidy between the two sexes by halving the expression – silencing of one X in female (Heard et al. 1997). Although dosage compensation is well studied in sex chromosome evolution, whether and how it evolves after a chromosome ploidy variation is unclear, particularly for autosomes (Kojima and Cimini 2019). Previous studies did not report consistent results, with a significant increase of expression for the duplicated genes in *C. albicans* (Selmecki et al. 2006), *Drosophila* (Loehlin and Carroll 2016), *S. cerevisiae* (Torres et al. 2007), mammalian cells (Williams et al. 2008); and a decrease of the expression of the two copies in yeast and mammals, suggesting a compensation at the transcriptional level (Henrichsen et al. 2009; Qian et al. 2010). In the case of chromosome duplication, evidence for dosage compensation at the transcriptional level is scarce (Stenberg et al. 2009; Hose et al. 2020), and scaling of gene expression with gene copy number has been reported in disomic yeasts (Kaya et al. 2020). Interestingly, dosage compensation has been reported to occur at the post-transcriptional level via the modification of translation efficiency in *Drosophila* (Zhang and Presgraves 2017) or at the post-translational level for 20% of the proteome in yeast via an increased degradation rate of proteins (Dephoure et al. 2014). Increase of protein degradation has also been reported to be involved in compensation mechanism in human Downs syndrome for proteins encoded on the triplicated chromosome 21 and include known stable heteromeric protein complexes (Liu et al. 2017).

Although the consequences of chromosome duplication on transcription rates - and to a lesser extent their consequences on translation rates and protein abundance - have been investigated in many model organisms, our knowledge of the rate of chromosome duplication events in eukaryotes have been yet restricted to *Saccharomyces cerevisiae* (Lynch et al. 2008; Zhu et al. 2014; Liu and Zhang 2019) and humans (Nagaoka et al. 2012; Loane et al. 2013). Here, we investigate the spontaneous chromosome duplication rate in six unicellular photosynthetic species by analyzing mutation accumulation (MA) experiment genomes. The principle of MA experiments is to follow the descendants originated from a single cell under minimal selection insured by serial bottlenecks during dozens to thousands cell divisions (Halligan and Keightley 2009). Here, MA experiments have been performed by maintaining 12 to 40 MA lines for a total of 1,595 to 17,250 generations depending of the species (Krasovec et al. 2017; Krasovec, Sanchez-Brosseau, et al. 2018; Krasovec et al. 2019). The six species include five Chlorophyta (one Trebouxiophyceae et four Mamiellophyceae) and one Bacillariophyta, ecological relevant primary producers in the sunlit ocean (de Vargas et al. 2015), with a large phylogenetic spread encompassing 1.5 billion years of divergence (Yoon et al. 2004). We further recovered cryopreserved MA lines to investigate the consequence of chromosome duplication on the transcription and translation rates.

## RESULTS

### Whole chromosome duplication rate

In this study, we analyzed whole genome resequencing data from previous MA experiments (Krasovec et al. 2016; Krasovec et al. 2017; Krasovec, Sanchez-Brosseau, et al. 2018) in five haploid green algae species (Chlorophyta, four Mamiellophyceae and one Trebouxiophyceae) and one diploid diatom (Bacillariophyta) : *Picochlorum costavermella* RCC4223 (Krasovec, Vancaester, et al. 2018), *Ostreococcus tauri* RCC4221 (Blanc-Mathieu et al. 2014), *O. mediterraneus* RCC2590 (Subirana et al. 2013), *Bathycoccus prasinos* RCC1105 (Moreau et al. 2012), *Micromonas commoda* RCC299 (Worden et al. 2009) and *Phaeodactylum tricornutum* RCC2967 (Bowler et al. 2008; Giguere et al. 2022). Coverage was used as a proxy of copy number (Figure 1A and Supplemental figures S1 to S16), and unveiled whole chromosome duplication events in four of the six species (Table 1 and Table S1). Four whole chromosome duplications were observed in *M. commoda* (chromosomes C05, C12, C16 and C17), four in *B. prasinos* (chromosomes C04, C05, C06 and C19), one in *O. mediterraneus* (chromosome C14) and four in *P. tricornutum* (chromosomes C02, C14 and C23). The chromosome duplication events were mapped onto the genealogies of the MA lines to identify all independent chromosome duplication events (Figure 1B for *B. prasinos*). Genealogy analysis provided evidence that several chromosomes were duplicated two times independently: the MA lines Bp25c and Bp28b carry two independent duplications of chromosome C19; Mc08 and Mc09 carry two independent duplications of chromosome C17; and Pt11 and Pt10c carry two independent duplications of chromosomes C23. The probability of observing two independent whole chromosome duplication of the same chromosome in *B. prasinos, M. commoda* and *P. trichornutum* are 0.46, 0.47 and 0.34 respectively (see methods), and these probabilities are thus consistent with an equal probability of duplication across chromosomes. All independent duplication events inferred from coverage analysis and genealogies are summarized in Table 1 and S1. Unexpectedly, the analyses revealed that the ancestral line of the MA experiment in *B. prasinos* carried two copies of the chromosome C01. One of the copies had subsequently been lost 11 times independently over 4,145 generations, corresponding to a spontaneous chromosome loss of a duplicated chromosome of 0.006 per duplicated chromosome per generation in *B. prasinos*.

**Table 1.** Spontaneous whole chromosome duplication rate in six species. *N*_*lines*_ : number of MA lines, Gen : average number of generations per MA line, *Tot*_*Gen*_ : total number of generations, *N*_*Chrom*_ : number of chromosomes in the ancestral karyotype, *N*_*WCD*_ is the number of independent whole chromosome duplications, *U*_*WCDl*_ is the whole chromosome duplication rate per chromosome per cell division, and *U*_*Cell*_ is the whole chromosome duplication rate per cell division, *U*_*bs*_ is the base substitution mutation rate per cell division. The estimation of the upper limit of *N*_*WCD*_ *and U*_*cell*_ in *O. tauri* and *P. costavermella* relies on the assumption of one duplication event.

| Species | $N_{lines}$ | Gen | $Tot_{Gen}$ | $N_{Chrom}$ | $N_{WCD}$ | Duplicated Chrom | $U_{WCD}$ | $U_{cell}$ | $U_{bs}$ |
| --- | --- | --- | --- | --- | --- | --- | --- | --- | --- |
| *Bathycoccus prasinus* | 35 | 265 | 4,145 | 19 | 5 | C04, C05, C06, C19 | 0.000063 | 0.00121 | 0.0046 |
| *Micromonas commoda* | 37 | 272 | 4,994 | 17 | 5 | C05, C12, C16, C17 | 0.000059 | 0.00100 | 0.0171 |
| *Ostreococcus tauri* | 40 | 512 | 17,250 | 20 | 0 | - | <0.000003 | <0.00006 | 0.0054 |
| *Ostreococcus mediterraneus* | 33 | 272 | 8,380 | 19 | 1 | C14 | 0.000006 | 0.00012 | 0.0064 |
| *Picochlorum costavermella* | 12 | 133 | 1,596 | 10 | 0 | - | <0.00006 | <0.00063 | 0.0119 |
| *Phaeodactylum tricornutum* | 36 | 181 | 6,516 | 25 | 4 | C02, C14, C23 | 0.000025 | 0.00061 | 0.0132 |

**Figure 1.**
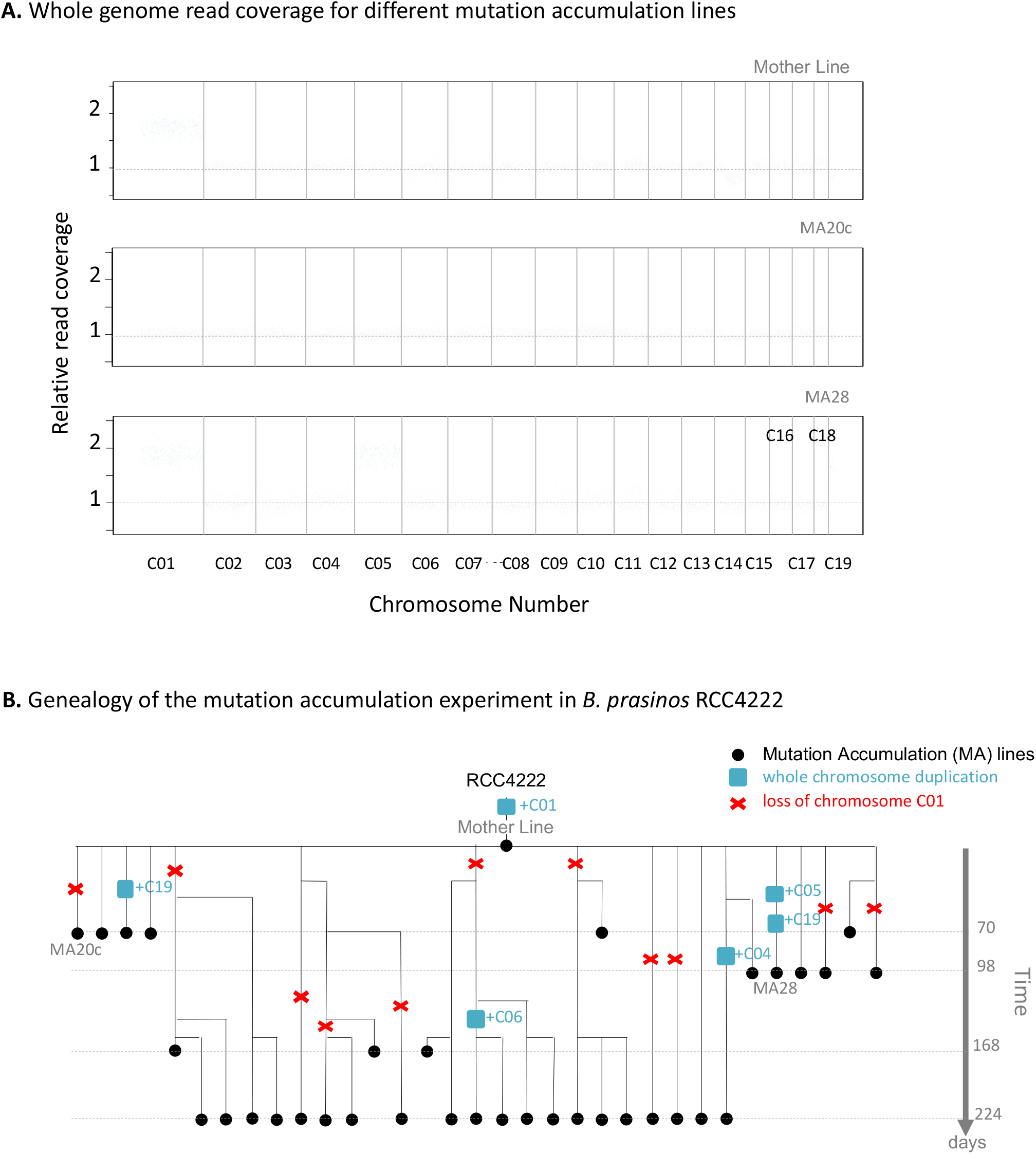
A. Normalized raw genomic coverage of the mother line (T0 of mutation accumulation experiment), Bp20c and Bp28 mutation accumulation lines of *Bathycoccus prasinos*. Grey lines are chromosome separators. In blue are the chromosomes in double copies. Raw coverage of all lines from all species are provided in Figures S1 to S16. **B**. Pedigree of the mutation accumulation lines from the *B. prasinos* experiment. Chromosome C01 is duplicated in the T0 line (named mother line) of the experiment. This duplication is then lost several times, and five other duplications of chromosomes C04, C05, C06 and C19 occurred.

### Consequences of chromosome duplication on transcription

To explore the consequences of the whole or partial chromosome duplication on transcription, we estimated the level of transcription of a control strain (*B. prasinos* RCC4222) and one MA line of *B. prasinos* by recovering one 4-year-old cryopreserved MA line. This cryopreserved culture originated from MA line Bp37, and the recovered culture is hereafter named Bp37B. Chromosome copy number of Bp37B and the control line were estimated by whole genome resequencing. This confirmed the two copies of chromosome C04 in Bp37B as in the original Bp37 (Figure 2A and S7), and also revealed additional karyotypic changes: an increase in chromosome C01 copy number (Figure 2A, Figure S17) in Bp37B. The heterogeneous coverage of chromosome C01 led us to divide it into two regions for subsequent analyses; region C01a (1.98 fold coverage) and C01b (1.35 fold coverage). The control line also contained duplicated regions (Figure 2A, Figure S17), chromosome C10 and a region of chromosome C02 named C02b, while C02a is in single copy. Limits of sub-regions of chromosomes C01 and C02 are provided in Table S2. We interpreted coverage values smaller than 2 (Figure S17), *e*.*g*. C01b in Bp37B, C10 and C02b as duplications that are carried by a subpopulation of cells. Cultures were grown to up to 10 million cells per ml prior to extraction so that polymorphism is not unexpected.

**Figure 2.**
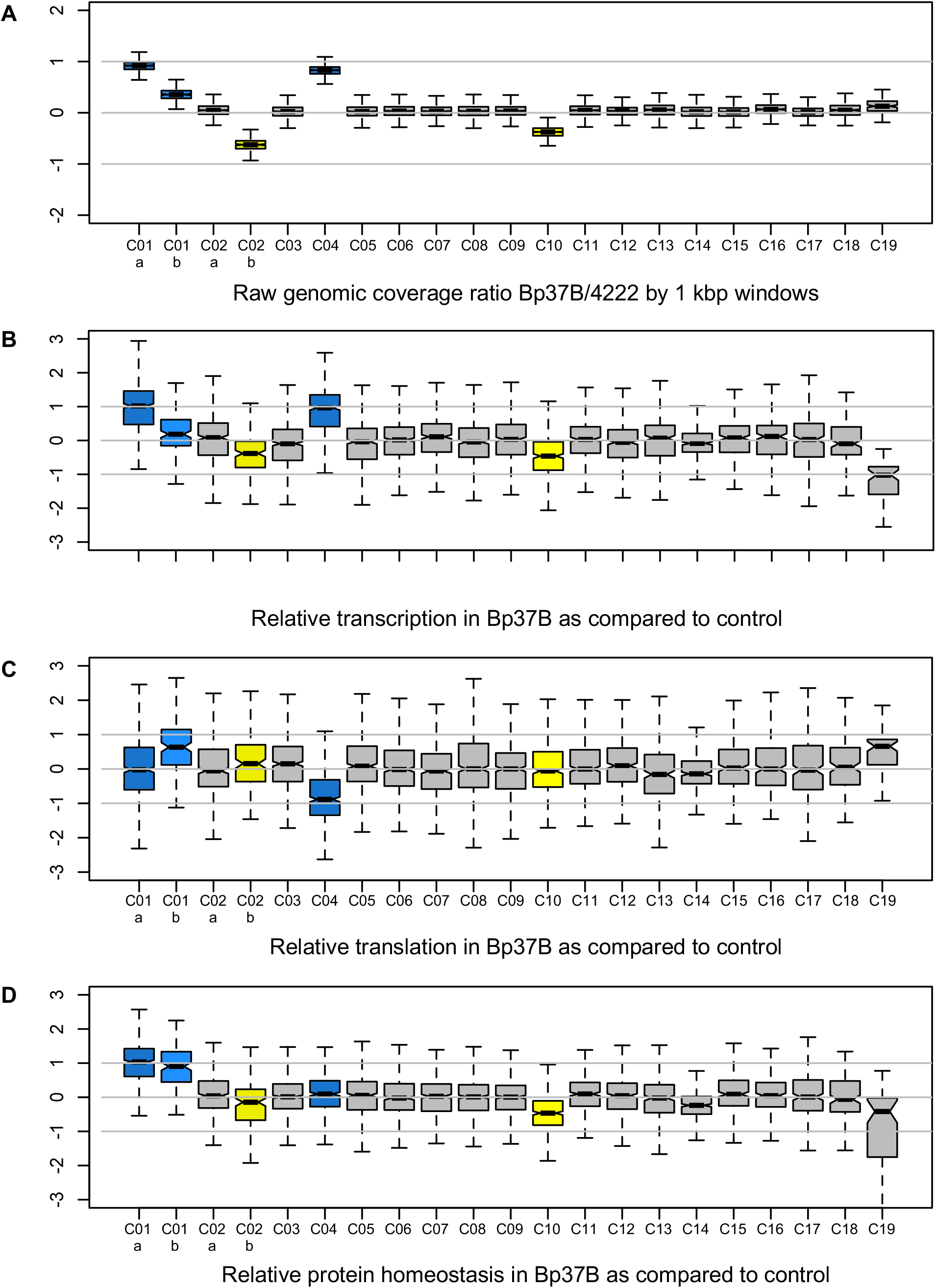
A. Genomic coverage of 1 kbp windows along the genome of the MA line *Bathycoccus prasinos* Bp37B. **B**. Distribution of genes transcription rate *tr(i)* of the 19 chromosomes. **C**. Distribution of genes translation efficiency *te(i)* of the 19 chromosomes. **B** and **C**. The transcription rate and translation efficiency ratio averages are different between chromosomes (ANOVA, p-value<10^−15^). **D**. Distribution of genes protein homeostasis (transcription rate x translation rate) of the 19 chromosomes. Y axis are in log2. Blue: duplications in Bp37B. Yellow: duplications in control RCC4222.

Comparative transcriptome analyses of Bp37B and the control line revealed that there are on average twice the number of transcripts for genes located on the duplicated chromosomes as compared to the control line - C04 (transcription rate *tr(i)* average=2.13, median=1.95, estimated from TPM, Figure 2B, raw data in Table S3) and C01a (transcription rate *tr(i)* average=2.29, median=2.02). The transcription rate of genes on chromosomes C10 and C02b is also affected. Altogether, the transcript production scales up with the chromosome copy number in the Bp37B and control lines (Pearson correlation, rho=0.87, p-value<0.001, Figure 3A).

**Figure 3.**
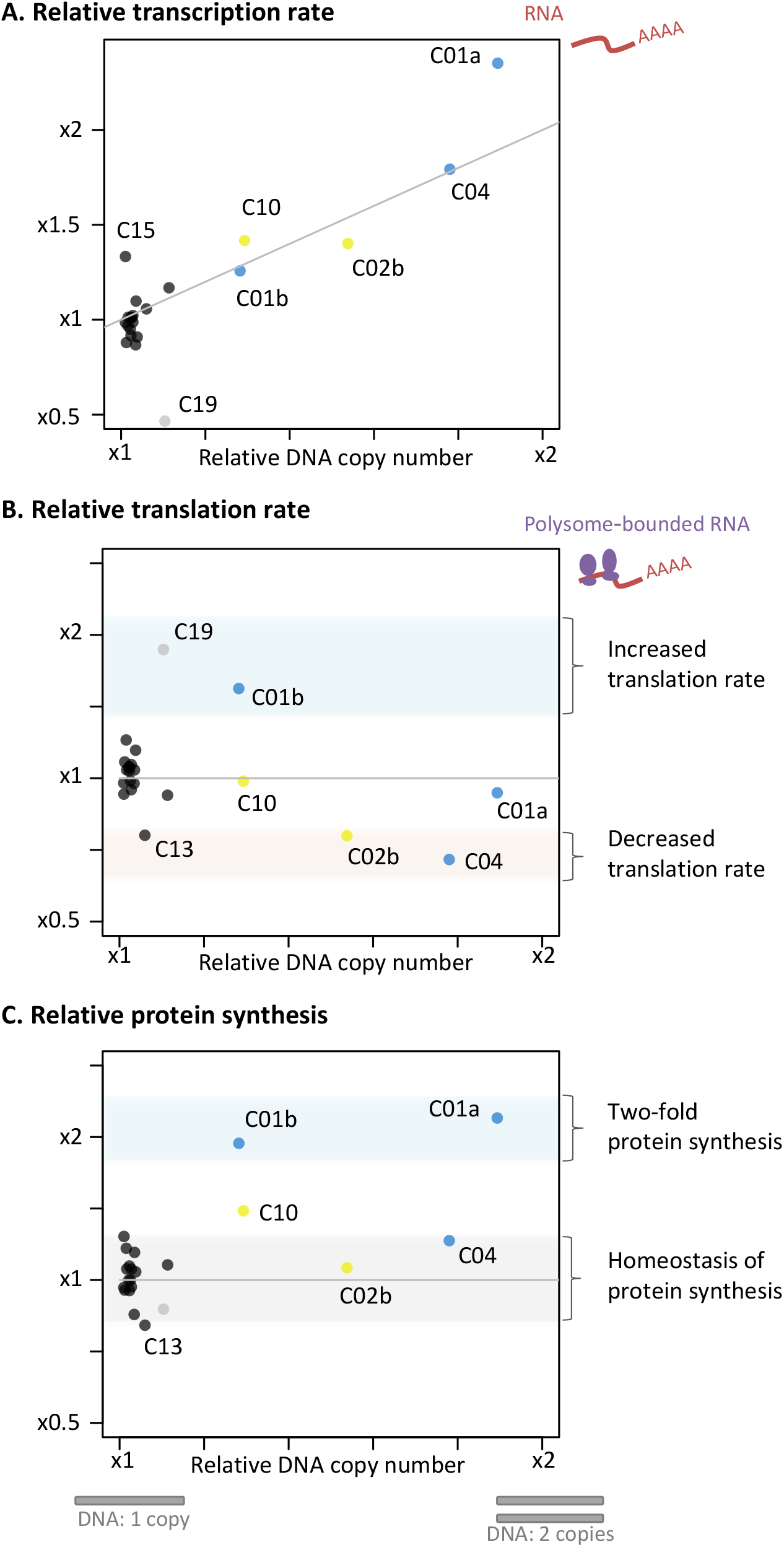
Raw relative genomic coverage average (Bp37B/Control) related to average transcription rate *tr(i)* (**A**), translation efficiency *te(i)* (**B**), and average protein synthesis (average transcription rate x average translation efficiency) per chromosome (**C**). Blue: chromosomes duplicated in Bp37B; Yellow: chromosomes duplicated the control RCCC4222; Black: non-duplicated chromosomes; Grey: outlier chromosome C19. Data are provided in Table S6 and S7.

### Gene-by-gene variation of the consequences of DNA copy number

Dosage invariant genes (Antonarakis et al. 2004; Lyle et al. 2004) are genes for which the transcription is not affected by copy number, and they may be involved in aneuploidy tolerance. We investigated differential gene expression at the individual gene scale with Deseq2 (Love et al. 2014) to identify candidate invariant genes. We found that the transcription rate of 158 out of the 1776 genes in two copies (9%) located on duplicated chromosomes or regions were not significantly higher than for genes in single copy (Table S4). Two Gene Ontology categories (RNA processing, GO:0006396 and DNA metabolic process GO:0006259) were over-represented in this transcriptomic invariant gene set. First, genes involved in RNA processing (3.8 times more frequent, p-value <0.01) including members of heteromeric protein complexes such as subunits 7 and 2 of the U6 snRNA-associated Sm-like protein (Table S5, invariant 158 annotations). Second, the genes involved in DNA metabolic process (2.2 times more frequent in the invariant gene subset, p-value <0.04) including the DNA Polymerase A and the DNA primase large subunit. Notable protein complex members of the invariant data set are Histone 3 and the subunit E of the translation initiation factor 3. There are in total 12 genes annotated as subunits in the subset of invariant genes, this is significantly more than the frequency of protein coding genes annotated as subunits in the complete proteome (Fisher exact test p=0.0006). This suggests that a higher proportion of these genes coding for protein forming complexes have a gene-specific tailored regulated transcription that is not affected by gene copy number.

### Consequences of chromosome duplication on translation

The compelling evidence of overexpression of the majority of duplicated genes prompted us to investigate whether post-transcriptional processes may temperate this excess of transcripts. This second hypothesis was tested in the same *B. prasinos* MA line by sequencing mRNAs associated with ribosomes (polysomes) in order to compare the translation efficiency *te(i)* of genes located on duplicated and non-duplicated lines, by estimating the relative proportion of mRNA binded to a ribosome in single versus duplicated genes. We found that the average translation efficiency of the genes located on the duplicated chromosome C04 was 0.71 (median=0.54) as compared to the genes on this chromosome in the control line (Figure 2C). At chromosome scale (Table S7), the translation rate is 0.67 for the chromosome C04. However, the translation efficiency of part C01a of the chromosome C01 was 0.93 and 1.54 for the part C01b. The translation efficiency of the two parts of the chromosome C02 are 1.04 and 0.76 for C02a and C02b, respectively, and 0.99 for chromosome C10. Translation rate modification is thus chromosome dependent (Figure 3): genes located on the C04 and C01b display an important variation in translation efficiency between the two lines. Last, we estimated the expected protein production in Bp37B as compared to the control line by going back to the absolute translation rate for each gene: that is the ratio of mRNA in polysomes in Bp37B as compared to the control (Figure 2D). This predicted that genes on C04 have a similar protein production rate in Bp37B and in the control line, despite a higher transcription rate as a consequence of the chromosome duplication, as well for the two parts of the chromosome C02 (Figure 2D). However, the two parts of the chromosome C01, despite different transcription and translation efficiency rates both leads to a protein level that is the double of the control line. The chromosome C10 also exhibits a lower protein content in Bp37B compared to control, meaning there is an excess of proteins from genes linked to this chromosome in the control line because of the duplication. Altogether, we observed there is no dosage compensation at the transcription level for any of the duplicated regions, but that dosage compensation may occur at the translation level on some duplicated regions and seems chromosome dependent (Figure 3).

### Searching for independent molecular signatures of translation modulation

The observed dosage compensation at the translation level on chromosome C04 suggests that a post-transcriptional mechanism occurs on duplicated transcripts. As the length of the poly(A) tail is a key feature of many cytoplasmic mRNAs and is known to regulate translation (Subtelny et al. 2014; Eichhorn et al. 2016; Lim et al. 2016), we tested if DC observed for the translation of transcripts linked to duplicated chromosome C04 is associated with a variation of poly(A) tail length. Using 3’RACE experiment, we measured the poly(A) tail length in Bp37B and control (Figure 4 and S18) of the transcripts of three genes located on chromosome C04 (04g00840, 04g01730 and 04g04360), one gene located on chromosome C08 (08g03470) and C01 gene located on chromosome C01a (01g01300). We systematically observed that the poly(A) tail length is reduced in Bp37B line compared to control line for transcripts located on chromosome C04 (Figure 4), whereas no difference is observed control (Figure S18).

**Figure 4.**
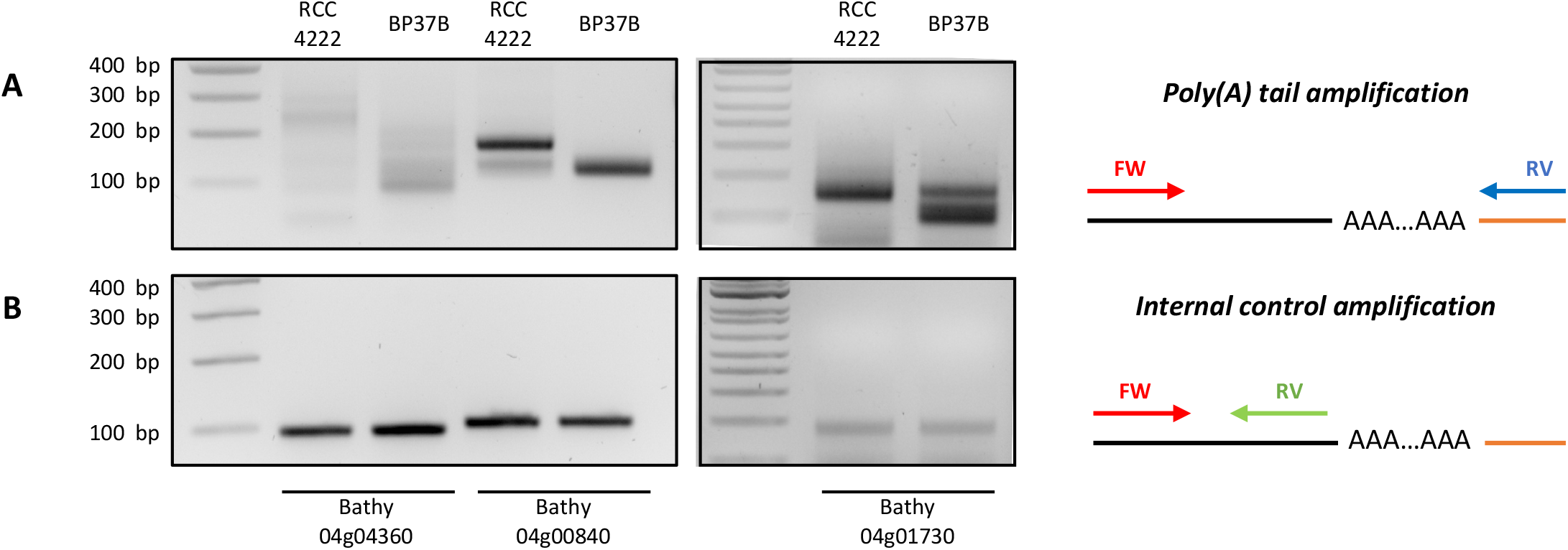
Transcripts from genes located on duplicated chromosome C04 present shorter poly(A) tail. Poly(A) tail measurement was performed using modified 3’RACE. PCR amplification was performed using primers flanking poly(A) tail (**A**) or primers anchored in 3’UTR just before poly(A) tail as internal control (**B**). Experiments was performed using total RNA from RCC4222 line (control line) and BP37B line (with duplicated chromosome C04). Illustrations representing PCR amplification are present on the right panel. Black line represents the messenger RNA and the orange line represent the ligated adapter used for reverse transcription and PCR amplification. FW. Forward primer. RV. reverse primer.

## DISCUSSION

### Rates of whole chromosome duplication in eukaryotes

The rates of spontaneous whole chromosome duplication reported here vary between 6×10^−5^ to 1×10^−3^ events per haploid genome per generation and are thus one to two orders of magnitude lower than spontaneous point mutations per genome per generation. Theoretically, there are two possible mechanisms at the origin of chromosome duplication i) the supernumerary replication of one chromosome before cell division, leading to one cell with one chromosome and one cell with two chromosomes; or ii) the unequal segregation of chromosomes during cell division, leading to one cell with two copies and one cell without any copy. The latter scenario has been intensively experimentally studied in human cells (Ford and Correll 1992; Cimini et al. 2001), and seems all the more likely in the extremely small-sized Mamiellophyceae cells (1 μm cell diameter) as it has been reported that they contain less kinetochore microtubules than chromosomes (Gan et al. 2011), which may induce high error rate in the chromosome segregation process. However, the whole chromosome duplication rates reported in Mamiellophyceae are in line with those found in the diatom *P. tricornutum* and in yeast (9.7×10^−5^) (Zhu et al. 2014), the latter containing approximately as many kinetochore microtubules as chromosomes (Peterson and Ris 1976). Altogether, the spontaneous whole chromosome duplication rates reported previously in yeast (Zhu et al. 2014) and in the species from evolutionary distant eukaryotic lineages reported here (Yoon et al. 2004) suggest a high rate of spontaneous aneuploidy across the eukaryotic tree of life. To investigate whether spontaneous whole chromosome duplication had been overlooked in other lineages, we screened publicly available resequencing data of MA on the freshwater green algae *Chlamydomonas reinhardtii* (Ness et al. 2015), and identified one event for chromosome 14 in the strain CC1373. We also provided evidence of spontaneous whole chromosome duplication (four chromosomes in a single individual from a pedigree) in the brown algae *Ectocarpus* (Krasovec et al. 2022).

Interestingly, we were also able to estimate the rate of chromosome duplication loss during one experiment to be about three orders of magnitude higher than the spontaneous duplication rate. This is consistent with previous observation in *S. cerevisiae*, and our observations that cells with duplicated chromosomes are ephemeral and unlikely to be maintained for a long time in batch culture. Indeed, the coverage of the large duplication events on C01b, C02b, C04 and C10 were 1.4,1.6, 1.8 and 1.4 times the coverage of single copy chromosomes respectively, suggesting that the proportion of cells carrying a supernumerary chromosome within the population is 0.4, 0.6 and 0.8. In addition, we observed the loss of the chromosome C04 duplication in one of the Bp37B lines over 4 months of sub-culturing.

As opposed to the aneuploidy generated during meiotic divisions, which has been intensively studied in mammalian cells (Nagaoka et al. 2012), the chromosomal duplication events reported here are generated during mitotic division. Since the molecular mechanisms involved in chromosomal segregation in mitosis, meiosis 1 and meiosis 2, are different, the associated aneuploidy rates are not expected to be equal. The aneuploidy rates in mitotically dividing human cells, such as HCT116 lines, has been estimated to be 7×10^−2^ (Thompson and Compton 2008), more than one order of magnitude higher than the highest estimation of 1.2×10^−3^ reported here in *B. prasinos*. The observed difference of the spontaneous point mutation rate per genome per generation is expected to vary between species as the consequence of different genome sizes, effective population sizes (Lynch et al. 2016), and the difference between the observed and expected GC content (Krasovec et al. 2017). However, the only eight spontaneous aneuploidy rates yet available are not sufficient to investigate the reasons of chromosome duplication rate variations, such as the impact of chromosome number or effective population size.

### Fitness effect of whole chromosome duplication and dosage compensation

The high spontaneous rates of spontaneous chromosome duplication challenge the common idea that whole chromosome duplications are highly deleterious for individual cells. Using the number of cell divisions per days as a proxy of fitness during the MA experiment (Krasovec et al. 2016), only one out of the 15 MA lines with a whole chromosome duplication displayed a significant fitness decrease (Bp28b, Pearson correlation, p-value=0.016, *ρ*=-0.846, Table S8). Recent studies in yeast are inconsistent with a highly deleterious fitness effect of aneuploidy and point towards slightly deleterious effects, as well as a significant effect of the genetic background on aneuploidy tolerance (Scopel et al. 2021). Compensating point mutations have been previously linked to aneuploidy tolerance in yeast (Torres et al. 2010), notably in the gene encoding the deubiquitinating enzyme UBP6. Point mutations are unlikely to impact the aneuploidy tolerance of the different MA lines reported here, as some MA lines with chromosome duplication do not carry any point mutation (Mc3, Bp28b, Bp26 and Bp25), or carry only synonymous mutations and mutations located in intergenic region (*i*.*e*. Om3 or Mc28, Table S9). The deleterious effect of aneuploidy could be mitigated by a dosage compensation mechanism preserving a relative protein homeostasis. Here in *B. prasinos*, despite the important variation in transcription and translation between genes, we observed that transcription rates overwhelmingly scaled up with chromosome copy number (Figure 3A) and that translation rates differed significantly between chromosomes (Figure 3B). However, chromosome C19 displays a unique pattern. This chromosome is a small hypervariable outlier chromosome found in all Mamiellophyceae species sequenced so far (Blanc-Mathieu et al. 2017; Yau et al. 2020). Previous studies reported a strong variation in transcription rates of genes on this chromosome, which is associated with resistance to viruses (Yau et al. 2016) and characterized by lower GC content than other chromosomes.

For the duplicated chromosomes or regions, we observed three different patterns of translation efficiency. First, we observed no relative difference in translation rates for genes on duplicated chromosome C10, which is duplicated in 40% percent of cells, and region C01a, which is duplicated in 100% of cells. The inferred excess of protein synthesis of duplicated genes on this chromosome is thus 40% (Figure 3C). Second, we observed a decrease of the translation rate for genes linked to chromosome C04 and region C02b. More precisely, the estimated percent of mRNA linked to ribosomes is inversely proportional to the excess of mRNA produced for duplicated genes on these two chromosomes, predicting approximately the same protein production rate as in cells without duplicated chromosomes (Figure 3C). Third, and very surprisingly, we observed an increase of the translation level of genes on C01b, that are in two copies in 40% of cells. As a consequence, the rate of protein synthesis on genes located on region C01b reach the same relative protein synthesis than genes located on region C01a, that is twice the protein synthesis predicted in cells with a single copy of this chromosome. We speculate the absence of dosage compensation on region C01a and the excess of translation on region C01b simulate the ancestral protein homeostasis in the mother line, which contained two entirely duplicated chromosome C01.

Chromosome-wide regulation of gene translation is relatively unexplored and poorly understood. Indeed, while location on a chromosome may predict a gene’s transcription rate as a result of epigenetic marks on the DNA, epitranscriptomics chromosome wide mRNA modifications mechanisms are yet to be discovered. Notwithstanding, chromosome wide translational dosage compensation has been previously observed in *Drosophila* (Zhang and Presgraves 2016), and at the gene scale in several species (Zhang and Presgraves 2017; Chang and Liao 2020).

### Poly(A) tail length

The role of poly(A) tail length in the translation efficiency is starting to be better understood (Weill et al. 2012; Subtelny et al. 2014), and alternative polyadenylation is indeed implicated in several processes: transcription termination by RNAP II, mRNA stability, mRNA export and translation efficiency (Zhang et al. 2010; Di Giammartino et al. 2011). In the cytoplasm, the poly(A) tail plays important roles in mRNA translation and stability and the modulation of its length has an important impact on translation efficiency (Subtelny et al. 2014; Eichhorn et al. 2016; Lim et al. 2016). As example, during oocyte maturation and early embryonic development, an increase of poly(A) tail length occurs for particular mRNAs resulting in an increase of translation (Subtelny et al. 2014; Eichhorn et al. 2016; Lim et al. 2016). This modulation of poly(A) tail length has also been found to activate some neuronal transcripts (Udagawa et al. 2012). Recently, it has been suggested that poly(A) tails can be modulated to balance mRNA levels and adjust translation efficiency (Slobodin et al. 2020). In this last case, the CCR4-Not complex shortened the poly(A) tails that reduces the stability of mRNAs. Here, our data suggest that, in a context of chromosome duplication, modulation of poly(A) tail length could be a key post-transcriptional mechanism necessary for dosage compensation at translation level. Future work to explore genome wide analysis of poly(A) tail variation and post-transcriptional RNA modifications such as adenosine methylation (Miao et al. 2022) are poised to clarify the translation compensation mechanisms.

## CONCLUSION

In conclusion, the high prevalence of whole chromosome duplication in five unicellular photosynthetic eukaryotes suggest spontaneous whole chromosome duplication is pervasive in eukaryotes. In one species, *B. prasinos*, we provide evidence that a whole duplication chromosome event is associated with dosage compensation at the post-transcriptional level which might involve the adjustment of poly(A) tail length. These results point to yet unknown post-transcriptional regulation mechanisms in DC of aneuploid karyotypes.

## MATERIALS AND METHODS

### Sequencing data from mutation accumulation (MA) experiments

Mutation accumulation (MA) experiments of the six species *Picochlorum costavermella* RCC4223 (Krasovec, Vancaester, et al. 2018), *Ostreococcus tauri* RCC4221 (Blanc-Mathieu et al. 2014), *O. mediterraneus* RCC2590 (Subirana et al. 2013; Yau et al. 2020), *Bathycoccus prasinos* RCC1105 (synonym to RCC4222) (Moreau et al. 2012), *Micromonas commoda* RCC299 (Worden et al. 2009) and *Phaeodactylum tricornutum* RCC2967 (Bowler et al. 2008; Giguere et al. 2022) were conducted with a flow cytometry protocol described previously for phytoplankton species in liquid medium (Krasovec et al. 2016). Briefly, a mutation accumulation experiment consists in following of MA lines that have evolved from a same cell (the ancestral line) during hundreds of generations. Relaxed selection pressure on spontaneous mutations is ensured by maintaining all MA lines at very low effective population sizes (6<Ne<8.5) throughout the experiment (Krasovec et al. 2016). As a consequence, MA experiments enable to estimate spontaneous mutation rates, excluding lethal mutations. Coupled with whole genome sequencing of ancestral and MA lines, this experiment enables the direct estimation of the spontaneous mutation rate of a species. Here, MA lines came from a single cell obtained by dilution serving as *T*_*0*_ culture (named the mother line, ML) and were maintained in 24-wells plates in L1 medium at 20 °C with a 16h-dark 8h-light life cycle. Single cell bottleneck by dilution were done each 14 days to have a low effective population size and limit selection. DNA of initial line (ML line) and final time of MA lines were extracted with chloroform protocol and sequencing done with Illumina HiSeq or MiSeq by GATC biotech (Germany). To detect duplications, raw reads were mapped against the reference genomes for each strain with bwa mem v0.7.12 (Li and Durbin 2010). Then, bam files were treated with samtools v1.3.1 (Li et al. 2009) and bedtools v 2-2.18.0 (Quinlan and Hall 2010) to extract the coverage.

### Statistical analyses

The probability of drawing one chromosome twice out of *k* WCD events chosen from *n* chromosomes, *P*(*j,n*), is equal to the ratio of the number of combinations of (*k*-1) out of *n* without replacement multiplied by *k*-1, which corresponds to the number of possible *k*-1 different chromosomes with one in two copies, out of *n*, divided by the number of combinations of *k* out of *n* with replacement (*k*-1)\**C*(*k*-1,*n*)/(*n*+*k*-1), which corresponds to the total number of possible subsamples of k chromosomes out of *n*.

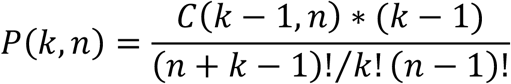

All statistical analyses were performed with R (R Core Team 2022).

### Dosage compensation analysis

This study was started three years after the end of the MA experiments, for which some MA lines had been cryopreserved which allowed us to re-start a culture of one *B. prasinos* MA line with a duplicated chromosome C04 (Bp37) to investigate possible dosage compensation. The restarted culture from the cryopreserved Bp37 was renamed Bp37B. As a control, we used the reference culture of the strain *B. prasinos* RCC4222, that is the derived from the RCC1105 used for the MA experiment. All cultures (Bp37B and RCC4222) were maintained under a 12:12 h light:dark regime under 50 μmol photon m^−2^ s^−1^ white light at 20 °C. The karyotype of the defrozen cultures Bp37B and the control RCC4222 were checked by DNAseq resequencing. The karyotype of Bp37B contained on additional chromosome copy number as compared to Bp37, and the karyotype of RCC4222 contained one additional copy numbers and one less chromosome copy number as compared to the mother line (ML) (Figure S17). The relative transcription rate was corrected by the number of DNA copy in each line for further analysis below.

For expression analysis, total RNA was extracted using the Direct-zol RNA MiniPrep Kit (Zymo Research, Californie, USA) from pooling flasks of cultures (100 ml cultures with 200 million cells per ml) taken 6h before and 1h before the light on, in triplicates for the control RCC4222 and duplicates for Bp37B. Then for translation efficiency analysis, polysome extraction was performed for Ribo-seq as described previously (Carpentier et al. 2020) with few modifications. Briefly, 600 mL of *B. prasinos* culture were centrifuged at 8,000 g for 20 minutes. After centrifugation, pellets were resuspended in 2.4 mL of polysome extraction buffer. After 10 minutes of incubation on ice and centrifugation, 2 mL of supernatant was loaded on a 9 mL 15–60% sucrose gradient and centrifuged for 3h at 38 000 rpm with rotor SW41 Ti. Fractions corresponding to polysomes were pooled and polysomal RNA was extracted as previously described (Carpentier et al. 2020). RNA library preparation was performed on total or polysomal RNA using a NEBNext Poly(A) mRNA Magnetic Isolation Module and a NEBNext Ultra II Directional RNA Library Prep Kit (New England Biolabs) according to the manufacturer’s instructions with 1 μg of RNA as a starting point. Libraries were multiplexed and sequenced on a NextSeq 550. Raw reads were mapped against the reference transcriptome of *B. prasinos* with RSEM with standard parameters (Li and Dewey 2011). We obtained the TPM average of total RNA and polysomes linked RNA that we compared between *B. prasinos* RCC4222 and the MA line Bp37B. At the gene scale, we first used the total RNA TPM values to estimate the expression difference ratio, *r*_*RNA*_ for each gene *i*, between two copies and single copy genes as:

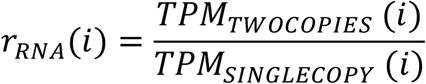

These relative expression rates *r*_*RNA*_*(i)* were normalized by the expression rate median of genes located on non-duplicated chromosomes in Bp37B (C03, C05, C06, C07, C08, C09, C011, C012, C013, C014, C015, C016, C017 and C018), in order to estimate the transcription rate *tr(i)* of duplicated chromosomes related to non-duplicated chromosomes. Chromosome C19 was not considered because it is a particularly outlier chromosome (see Discussion).

Second, we used the total RNA and polysomes linked RNA TPM values for each gene *i* to calculate the translation rates of each gene, *r(i)*, as:

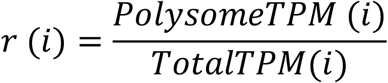

The translation efficiency between a gene *i* in the two lines in single versus two copies, *te(i)* was estimated as:

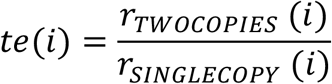

The *te*(*i*) ratio was then normalized by the median translation efficiency of all genes on non-duplicated chromosomes in strain BP37B (5,267 genes).

Then, to estimate the dosage compensation at the scale of the chromosomes, the values were normalized by the average TPM of all single copy chromosomes prior to the calculation of the ratios (Table S7).

Poly(A) tail analysis was performed as previously described with slight modifications (Sement and Gagliardi 2014). PCR products were resolved on a 2.5% agarose gel. Primers used in this study are available in Table S10.

### Differential gene expression analyses

The statistical significance of the genes differential expression levels was further estimated with Deseq2 (Love et al. 2014). Transcriptional invariant genes were defined as genes present on duplicated regions and for which the relative transcription rate *r*_*RNA*_*(i)* was comprised between 0.9 and 1.1. The over-representation of a certain GO term in the transcriptional invariant gene set was compared to the genome-wide GO term background frequency using the GO enrichment analysis with default values implemented in pico-PLAZA workbench (Vandepoele et al. 2013).

## AUTHOR CONTRIBUTIONS

MK, SSB and GP performed the mutation accumulation experiments. MK performed the bioinformatics analysis. FS performed the cell cultures, RNA extractions, PCR and gel migration experiments. RM designed and performed the polysome analyses, poly(A) tail tests and RNAseq preparation. GP performed statistical analyses and coordinated the project. MK drafted the first version of the manuscript and all authors participated to writing the final version.

## ACKNOWLEDGEMENTS

We are grateful to Claire Hemon and Elodie Desgranges for technical assistance with the MA experiments. We acknowledge the GenoToul Bioinformatics platform (Toulouse, France) for bioinformatics analysis support and cluster availability, the BIOPIC platform for support with the cytometry, and the sequencing facility of the Université de Perpignan Via Domitia BioEnvironnement platform. This work was funded by ANRJCJC-SVSE6-2013-0005 to GP and SSB and ANR PHYTOMICS (ANR-21-CE02-0026). This study is set within the framework of the “Laboratoires d’Excellences (LABEX)” TULIP (ANR-10-LABX-41) and of the “École Universitaire de Recherche (EUR)” TULIP-GS (ANR-18-EURE-0019).

## COMPETING INTERESTS

The authors declare no conflicts of interest.

## DATA AVAILABILITY STATEMENT

All genomic raw reads of ML and mutation accumulation lines are available under the bioprojects PRJNA531882 (*Ostreoccocus tauri, O. mediterraneus, Micromonas commoda, Bathycoccus prasinos*), PRJNA453760 and PRJNA389600 (*Picochlorum costavermella*), and PRJNA478011 (*Phaeodactylum tricornutum*). Transcriptomic raw reads of the *Bathycoccus prasinos* MA lines 37 (Bp37B) and control line RCC4222 are available under the bioproject PRJNA715163. A summary is provided in Table S11.

## SUPPLEMENTARY MATERIAL

**Figure S1 to S16**. Raw coverage by 1 kb windows of all mutation accumulation lines from *Ostreoccocus tauri, O. mediterraneus, Bathycoccus prasinos, Micromonas commoda, Picochlorum costavermella* and *Phaeodactylum tricornutum*. The horizontal grey lines indicate the chromosome separation, and the duplicated chromosome are in blue. ML is the T0 genome of mutation accumulation experiments. In *M. commoda*, a duplication of a fraction of chromosome 2 in the mother line (ML) is maintained in all MA lines, suggesting there might be a miss-assembly due to a duplicated region at this location.

**Figure S17.** Coverage analysis of the Bp37B and control RCC4222.

**Figure S18.** Transcripts from duplicated chromosome C01 and non-duplicated chromosome C08 present similar poly(A) tail.

**Table S1**. List of duplicated chromosomes in the MA lines.

**Table S2**. Limits of sub-regions of chromosomes C01 and C02.

**Table S3**. Raw transcript per million (TPM) table for all 10 samples.

**Table S4**. Estimation of the number of dosage insensitive genes per chromosome.

**Table S5**. List of the dosage insensitive genes with annotations.

**Table S6**. Relative copy numbers of chromosomes in the two lines of *Bathycoccus prasinos*.

**Table S7**. Relative TPM average per chromosome used for Figure 2.

**Table S8**. Fitness of MA lines with duplicated chromosomes during MA experiments. **Table S9**. Point mutations previously identified in the MA lines with whole chromosome duplications.

**Table S10**. Sequences of primers used for poly(A) tail length analysis.

**Table S11**. Summary of all accessions of the data used in this study with Bioprojects and biosamples.

---

**Figure S17.**
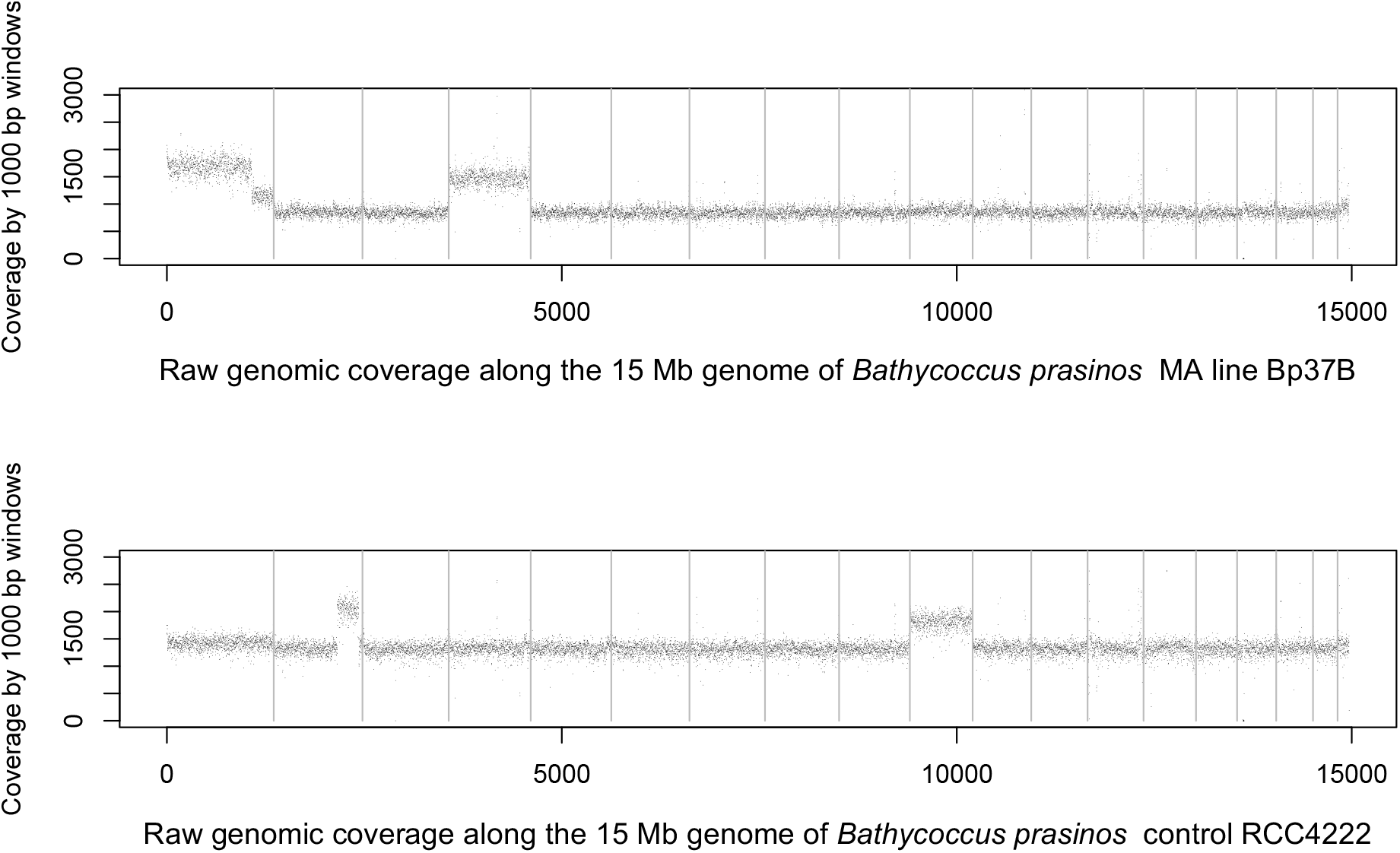
Coverage by 1 kb windows of the *Bathycoccus prasinos* MA line Bp37 and control RCC4222.

**Figure S18.**
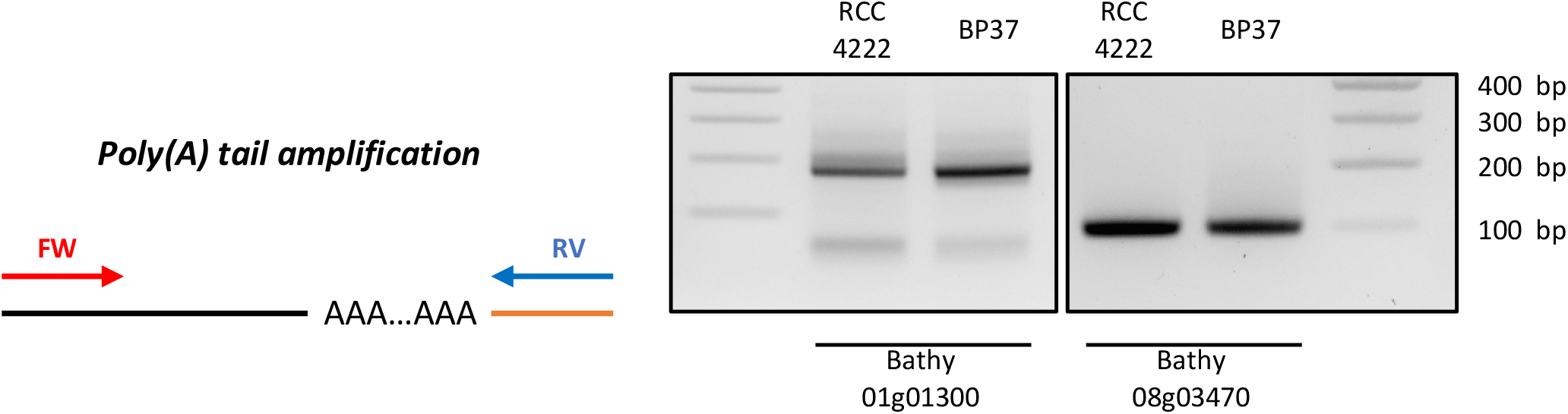
Transcripts from duplicated chromosome C01 and non-duplicated chromosome C08 present similar poly(A) tail. Poly(A) tail measurement was performed using modified 3’RACE. PCR amplification was performed using primers flanking poly(A) tail. Experiments was performed using total RNA from RCC4222 line (control line) and Bp37B line (chromosome C04 duplicated). Illustrations representing PCR amplification are present on the left panel. Black line represents the mRNA and the orange line represent the ligated adapter used for reverse transcription and PCR amplification.

## Notes

### Competing Interest Statement

The authors have declared no competing interest.

### Summary of Updates

new figures, additional analysis, re-writing of introduction and discussion

